# End-to-end assessment of fecal bacteriome analysis: from sample processing to DNA sequencing and bioinformatics results

**DOI:** 10.1101/646349

**Authors:** Ana Paula Christoff, Giuliano Netto Flores Cruz, Aline Fernanda Rodrigues Sereia, Laís EikoYamanaka, Paola Paz Silveira, Luiz Felipe Valter Oliveira

## Abstract

Intestinal microbiome, comprising the whole microbiota, their genes and genomes living in the human gut have significant roles in promoting health or disease status. As many studies showed so far, identifying the bacterial components of the microbiome can reveal important biomarkers to help in the disease comprehension to a further adequate treatment. However, the human nature is quite variable considering the genetic components associated with life styles, directly reflecting on the gut microbiome. Thus, it is extremely important to know the populational microbiome background in order to draw conclusions regarding the health and disease conditions. Also, methodological best practices and knowledge about the methods being used are essential for the results quality and applicability with clinical relevance. In this way, we standardized the sample collection and processing methods used for the Probiome assay, a test developed to identify the Brazilian bacteriome from stool samples. EncodeTools Metabarcode pipeline of analysis was developed to obtain the best result from the samples. This pipeline uses the information of amplicon single variants (ASVs) in 100% identical oligotype clusters, and performs a *de novo* taxonomical assignment based on similarity for unknown sequences. To better comprehend the results obtained in Probiome assays, is essential to know the intestinal bacteriome diversity of Brazilians. Thus, we applied the standardized methods herein developed and began characterizing our populational data to allow a better understanding of the Brazilian bacteriome profiles and how they can be related to other microbiome studies.

## Introduction

The human body hosts a diverse microbial community composed by bacteria, fungi, virus, and small eukaryotes that along with their genes and genomes comprise the human microbiome. All this microbial living in our bodies, mainly in our intestine, serves as a source of genetic and metabolic diversity. Most of our gut microbiota is composed of bacteria [1,2] and their diversity influence the human health by playing a role in the digestive, neurological, or immunological systems disorders [3–5]. Two larger projects made significant contributions in the understanding of the healthy microbiota and their host, the Metagenomics of the Human Intestinal Tract (MetaHIT) [1] and the Human Microbiome Project (HMP) [6,7]. More recently, the American Gut project also contributed to the knowledge of intestinal microbiome profiles from populations in the United States, United Kingdom and Australia [8]. These microbiome projects, along with several others conducted around the world, have the primary goal of understanding the dynamics and variations in the human intestinal microbiome to characterize it regarding health and disease conditions.

The intestinal microbiome varies widely among individuals, also fluctuating over human development and time. These variations increase the complexity of the human microbiome comprehension, becoming more challenging to define what is a healthy status for a population and an individual [9]. Additionally, each population has its particularities regarding their genetic background, physiology, lifestyle, nutrition, and habits that can influence the microbiota [10,11]. A recent study published with Chinese populations revealed that geography has a substantial interference with microbiome profiles, hampering the universal application of microbiota-associated disease models that were developed based on specific populations [12]. Thus, it is extremely relevant to have microbiome information about the specific target population to allow conclusions regarding their health and disease conditions.

All these research studies were fundamental to improve the knowledge regarding microbiome characterization along with the technical and biological challenges that must be addressed and controlled in the best possible ways [13]. The experimental reproducibility is critical, giving the potential of clinical application for the obtained results. Moreover, adequate sample collection and storage is a requirement for maintaining the original microbial composition, since the improper storage can allow selective microorganisms to overgrow leading to microbial profile biases and consequently misleading the interpretation of the results [14,15]. Several efforts have also been made to address variations and standardize DNA extraction, amplicon 16S rRNA gene sequencing, and bioinformatics analysis, as done by the Microbiome Quality Control (MBQC) project consortium [16]. Moreover, usage of amplicon sequence variants (ASVs), the exact DNA sequence read, instead of the OTU picking (generally clustering sequences at 97% similarity) improves the resolution for microbiome results [13,17,18].

In this paper, we present an end-to-end assessment of a human intestinal bacteriome analysis for Brazilian populations, covering all the process from sample storage, amplicon library preparation, high-throughput DNA sequencing, and bioinformatics analysis. We introduced a new pipeline of analysis: EncodeTools Metabarcode, and generated 16S rRNA amplicon data for fecal samples of the Brazilian subjects to begin an understanding of the bacteriome compositional patterns in such a diverse population whose gut microbiome profiles are yet to be characterized.

## Material and Methods

### Sample collection and processing

Stool samples were collected using the Probiome kit (BiomeHub, Brazil) which includes a sanitary seat cover capable of retaining the stool and allows the proper sample collection with a sterile flocked swab - 520CS01 (Copan, USA) or 25-3606-H BT (Puritan, USA). The swabs have a breakpoint that allows the swab tip containing the collected sample to be inserted into a provided microtube with 1ml of fecal stabilization solution - ZSample (BiomeHub, Brazil). Each subject can take the entire kit home and perform the fecal sample collection individually. The samples were homogenized by microtube inversion and then forwarded to BiomeHub laboratory (Florianopolis, Brazil) for sample processing within 30 days after collection. In the laboratory, DNA was extracted from the preserved stool using the DNeasy PowerSoil Kit (QIAGEN, Germany) according to the manufacturer instructions. At each batch of DNA extraction, a negative control was included (CNE). A set of 206 stool samples that used the above collection and processing methods were randomly selected from the mischaracterized BiomeHub database. No possible correlations or associations with the fecal donors can be made from this bacterial sequences or any data included in this study. These samples, collected and anonymously processed as described above, along 2018, represent a Brazilian populational diverse subset comprising 65.4% female and 34.5% male from various geographical locations.

### Experimental subsets for sample storage, ZSample stability and DNA extraction tests

ZSample stability solution and stool sample preservation at room temperature were evaluated along 30 days. A single stool specimen was self-collected by an anonymous donor in seventeen replicates and stored in ZSample Probiome tubes to be analyzed at T0 (maximum of two hours after collection), T15 (15 days after sample collection) and T30 (30 days after collection). Five of the replicates were analyzed in T0, and six replicates were analyzed at each T15 and T30.

Additionally, batch effects for the ZSample lot production in stool sample preservation was evaluated along four batches of the solution produced at 0, 2, 9 and 18 months before the stool sample collection. Twenty-four replicates of a fecal sample from an anonymous donor were collected using the four solution lots listed above. For each lot, six replicates were obtained, three of them were processed in T0 (maximum of two hours after collection) and the other three in T30 (30 days after collection). All samples remained at room temperature in ZSample solution during the 30-day storage. Furthermore, these fecal samples collected and stored in ZSample were inoculated in a general culture media (PCA - plate count agar) and incubated at 35°C for three days, to evaluate cellular bacterial viability.

DNA extraction of fecal samples stored in ZSample was further tested in four different methods: DNeasy PowerSoil kit, DNeasy PowerSoil Pro kit, DNeasy PowerSoil Pro modified and QIAamp PowerFecal DNA kit, all from QIAGEN, Germany. In DNeasy PowerSoil Pro modified its original bead beating tubes with zirconium beads were replaced for the traditional PowerSoil silica bead tubes. Fecal samples were donated by five anonymous subjects, and processed with four experimental replicates for each extraction kit, in a total of 80 samples.

### DNA library preparation and sequencing

The *16S rRNA* amplicon sequencing libraries were prepared using the V3/V4 primers (341F CCTACGGGRSGCAGCAG and 806R GGACTACHVGGGTWTCTAAT) [19,20] in a two-step PCR protocol. The first PCR was performed with V3/V4 universal primers containing a partial Illumina adaptor, based on TruSeq structure adapter (Illumina, USA) that allows a second PCR with the indexing sequences similar to procedures described previously [21]. Here, we add unique dual-indexes per sample in the second PCR. Two microliters of individual stool sample DNA were used as input in each first PCR reaction. The PCR reactions were carried out using Platinum Taq (Invitrogen, USA) with the conditions: 95°C for 5 min, 25 cycles of 95°C for 45s, 55°C for 30s and 72°C for 45s and a final extension of 72°C for 2 min for PCR 1. In PCR 2 the conditions were 95°C for 5 min, 10 cycles of 95°C for 45s, 66°C for 30s and 72°C for 45s and a final extension of 72°C for 2 min. All PCR reactions were performed in triplicates. The final PCR reactions were cleaned up using AMPureXP beads (Beckman Coulter, USA) and samples were pooled in the sequencing libraries for quantification. At each batch of PCR, a negative reaction control was included (CNR). The DNA concentration of the libraries was estimated with Picogreen dsDNA assays (Invitrogen, USA), and then the pooled libraries were diluted for accurate qPCR quantification using KAPA Library Quantification Kit for Illumina platforms (KAPA Biosystems, MA). The libraries pools were adjusted to a final concentration of 11.5 pM (for V2 kits) or 18 pM (for V3 kits) and sequenced in a MiSeq system (Illumina, USA), using the standard Illumina primers provided in the manufacturer kit. Single-end 300 cycle runs were performed using V2x300, V2x300 Micro, V2x500 or V3x600 sequencing kits (Illumina, USA), always generating 283bp size amplicons suitable for analysis. Coverage of 50,000 reads was set to each sample sequenced.

### Bioinformatics analysis - EncodeTools Metabarcode pipeline

The sequenced reads obtained were processed using EncodeTools Metabarcode pipeline (BiomeHub, Brazil) a bioinformatics pipeline developed *in-house* and described below. Illumina FASTQ files were quality filtered and the primers were trimmed to yield a resulting read of 283bp. Only one mismatch is allowed in the primer sequences and the whole read is discarded if this criterion is not met. Sequenced reads smaller than expected or with remaining Illumina sequence adapter were discarded. After this initial quality assessment, identical read sequences (100% identity) were grouped into oligotypes and analyzed with Deblur package [22] to remove possible erroneous reads. After, VSEARCH [23] was used to remove chimeric amplicon reads. The oligotype clusterization with 100% identity provides a higher resolution for the amplicon sequencing variants (ASVs), also called sub-OTUs (sOTUs) [13] - herein denoted as oligotypes. An additional filter was implemented to remove oligotypes below the frequency cutoff of 0.2% in the final sample counts, *i.e.*, given a library size of 1,000 reads, oligotypes with less than two reads were filtered out. We also implemented a negative control filter, as in each processing batch we have negative controls for the DNA extraction and PCR. If any oligotypes were observed in the negative controls, they are checked against the samples and automatically removed from the sample results if present. The remaining oligotypes in the samples were used for taxonomic assignment with the BLAST tool [24] against a reference genome database. This database was constructed with complete and draft bacterial genomes, focused on clinically relevant bacteria, obtained from NCBI and *in-house* genome sequencings. It is composed of 11,750 sequences comprising 1,843 different bacterial taxonomies. Taxonomy was assigned to each oligotype using a lowest common ancestor (LCA) algorithm. If more than one reference can be assigned to the same oligotype with equivalent similarity and coverage metrics (*e.g.* two distinct species mapped to oligotype “A” with 100% identity and 100% coverage), the EncodeTools Metabarcode Taxonomy Assignment algorithm leads the taxonomy to the lowest level of possible unambiguous resolution (genus, family, order, class, phylum or kingdom), according to the similarity thresholds previously established previously [25]. The bacterial profile obtained at the end of the pipeline is shown in taxonomy proportions for the analyzed sample.

### Experimental subsets for robustness, sensibility and specificity of the EncodeTools Metabarcode pipeline

EncodeTools Metabarcode pipeline was tested and calibrated using internal data generated on diverse hospital microbiome DNA samples obtained and processed as previously described [26]. Eight different microbiome samples were evaluated (A-H). Seven of them (A-G) were diverse environmental swab samples and one was an artificial microbial community - mock (sample-H) - composed of: *Acinetobacter baumanii, Bacillus subtilis, Enterococcus faecalis, Escherichia coli, Klebsiella pneumoniae, Listeria monocytogenes, Pseudomonas aeruginosa, Salmonella enterica* and *Staphylococcus aureus.* The 16S rRNA amplicon library preparation for these eight different samples (A-H) was processed as described above in a total of 28 replicates per sample. These libraries replicates were prepared by three different operators in three separated MiSeq runs, totalizing 224 sample assays along with 22 negative controls. Eleven amplicon library replicates were prepared for each of the eight samples by a single operator for an intra-run technical reproducibility test and sequenced in a single V2x300 Illumina MiSeq run. Inter-run technical reproducibility test was done re-sequencing these eleven replicates amplicon libraries in a V3x600 Illumina MiSeq run. All sequencing runs were a single-end of 300 cycles. Then, two additional operators prepared the same amplicon libraries for the eight samples, in triplicates, for inter-run repeatability and robustness. These libraries were sequenced in two separated V2x300 Illumina MiSeq runs, one for each operator's library. All data generated were compared and used to evaluate the reproducibility, repeatability, sensibility and specificity for our amplicon library preparation along with DNA sequencing, and the EncodeTools Metabarcode pipeline of analysis.

### Data comparison and diversity analysis

The results from all samples were integrated into an oligotype table (analogous to OTU table), whose rows are samples and columns are oligotypes. For each oligotype, taxonomic lineage was computed. A typical data analysis input was comprised of oligotype, taxonomy, and metadata tables. The raw sequences were used to construct phylogenetic trees using FastTree 2.1 [27] and these were used to calculate weighted UniFrac [28] distances when suitable. Further analyzes were conducted inside the R statistical software environment (R version 3.6.0), using the Phyloseq package [29]. DESeq2, EdgeR, and metagenomeSeq packages were used for differential abundance analyses [30–32]. Nonparametric comparisons included *Kruskal-Wallis* and *Wilcoxon* tests as implemented in base R and in coin R package, respectively [33]. Other R packages used in this study are listed in Supplementary Table 1.

Alpha-diversity was computed using the plot_richness function from the Phyloseq R package with default parameters. Note that Phyloseq by default calculates the Simpson Diversity Index as 1 - D. Here, we transform the value back to 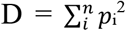 (*p*_*i*_ is the proportional abundance for the *i*^*th*^ taxonomy). Beta-diversity used proportion-normalized abundances as noted by [34] and [35]. Bray-Curtis Dissimilarity and weighted UniFrac were both calculated using Phyloseq’s distance function. Correlation coefficients between sample groups used mean taxonomy proportions within each group.

Differential abundance analysis was performed using four distinct methods, all of which using the above cited packages with default options unless stated otherwise: DESeq2 and EdgeR were used to fit Negative Binomial models with relative log expression scaling [30,31,35]; metagenomeSeq applied a zero-inflated log-normal model with cumulative-sum scaling [32]; finally, rarefaction (with Phyloseq) was also applied followed by exact *Wilcoxon-Mann-Whitney* test, as implemented in the Coin R package, as this is a very traditional method, even though it has been characterized by its lack of power [34,35]. Rather than accepting the significance calls from all methods or arbitrarily choosing one of them, here we considered as significantly differentiated those taxa that were detected by at least two distinct methods simultaneously. Effect sizes were reported as fold-changes in the *log*_2_ scale (*log*_2_ *FC*) for all but the *Wilcoxon-Mann-Whitney* method, whose effect size estimates were computed as 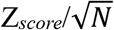 for sample size *N*. P-value correction for multiple comparisons was performed using the Benjamini-Hochberg procedure.

## Results

### Stool sample storage for bacteriome analysis

To validate our ZSample storage solution concerning the bacterial composition maintenance in fecal samples, we analyzed replicated samples stored at T0, T15 and T30 days. After DNA extraction and amplicon sequencing, we evaluated the bacterial profile from these samples through diversity and correlation analysis. Alpha and beta diversities showed no significant differences for the bacterial genera detected across sample storage times (Figures 1A and 1B). Additionally, high correlations were observed among bacterial profiles from all time points (Pearson and spearman’s > 0.92) (Figure 1C). Figures 1D and 1E show the bacterial relative abundance profiles across the replicated samples analyzed in T0, T15 and T30. Some variations could be observed; however, they were no more related to the storage time than with inter-replicates variation. The overall diversity and relative abundance of each bacterial genus detected remained equivalent in all the samples across the storage time. Data for correlations and bacterial abundance for other taxonomy levels (phylum, family and species) can be seen in Supplementary Figure 1. Taken together, these results indicate that ZSample properly maintains the original bacterial profile in samples stored at room temperature for at least 30 days. Moreover, no bacterial cellular viability was detected in the sample cultivation tests that were performed aerobically to resemble more closely how the samples are stored and manipulated along with the processes.

**Figure 1.**
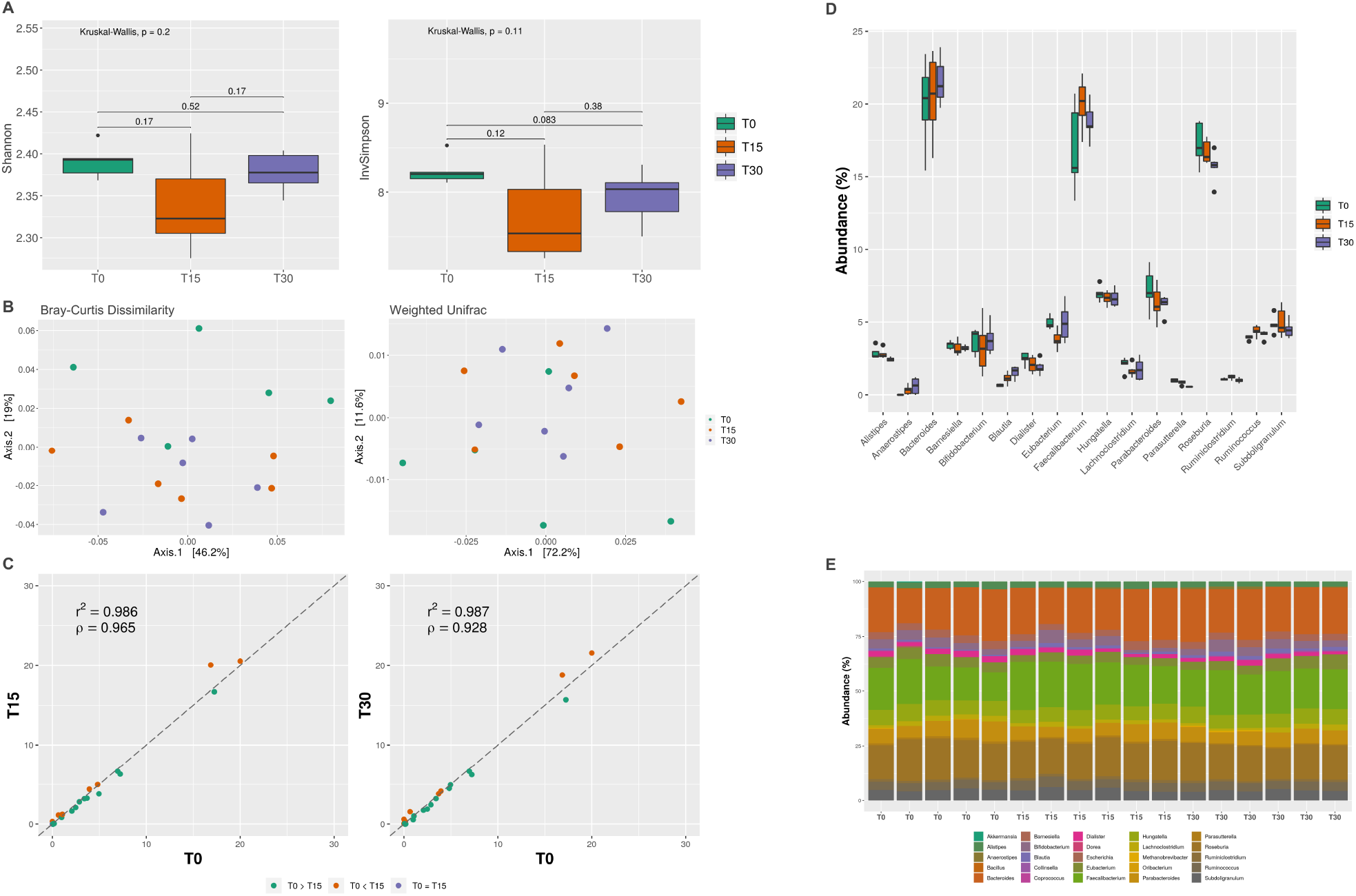
Fecal sample storage and bacterial profile along 30 days. Data presented in this figure is for bacterial genera analyzed after T0, T15 and T30 storage days in ZSample. **(A)** Shannon and InvSimpson alpha diversity analysis were performed with no significant differences among T0, T15 and T30 (*Kruskal-Wallis* p>0.05). Wilcoxon lacks of significance (p>0.05) is also showed above boxplots for pairwise comparisons between T0xT15, T15xT30 and T0xT30. **(B)** Beta diversities (Bray-curtis and Weighted UniFrac) didn’t show any specific sample grouping or deviation related to the storage time. **(C)** A correlation analysis was performed between T0-T15 and T0-30 showing values > 0.92 for Pearson (r^2^) and Spearman (ρ) coefficients. **(D)** Genera abundances along the sample storage demonstrate some inter-replicate variations higher than the storage time variation itself. **(E)** The proportional abundances for genera detected along the storage time in each replicate are shown. This also demonstrates the process reproducibility along different replicates and time.

As an additional validation step, we evaluated the batch effects of different ZSample lot productions in the bacterial profiles obtained in T0 or after 30 days (T30) of room temperature storage. High correlations (Pearson and Spearman’s > 0.94) were obtained for bacterial genera comparisons between storage in T0 and T30 (Figure 2A), and also for lots produced with differences in fabrication date of up to 18 months (Pearson and Spearman’s > 0.89) (Figure 2B). More detailed correlations considering other taxonomy levels as phylum, family and species are shown in Supplementary Figure 2. No significant of bacterial gain or loss due to the storage was observed in the data analyzed. Although relative abundances for bacterial phylum, family, genus or species demonstrate that the bacterial profile in the samples have some replicate variations, these were not correlated with the ZSampe production batch (Figure 2C).

**Figure 2.**
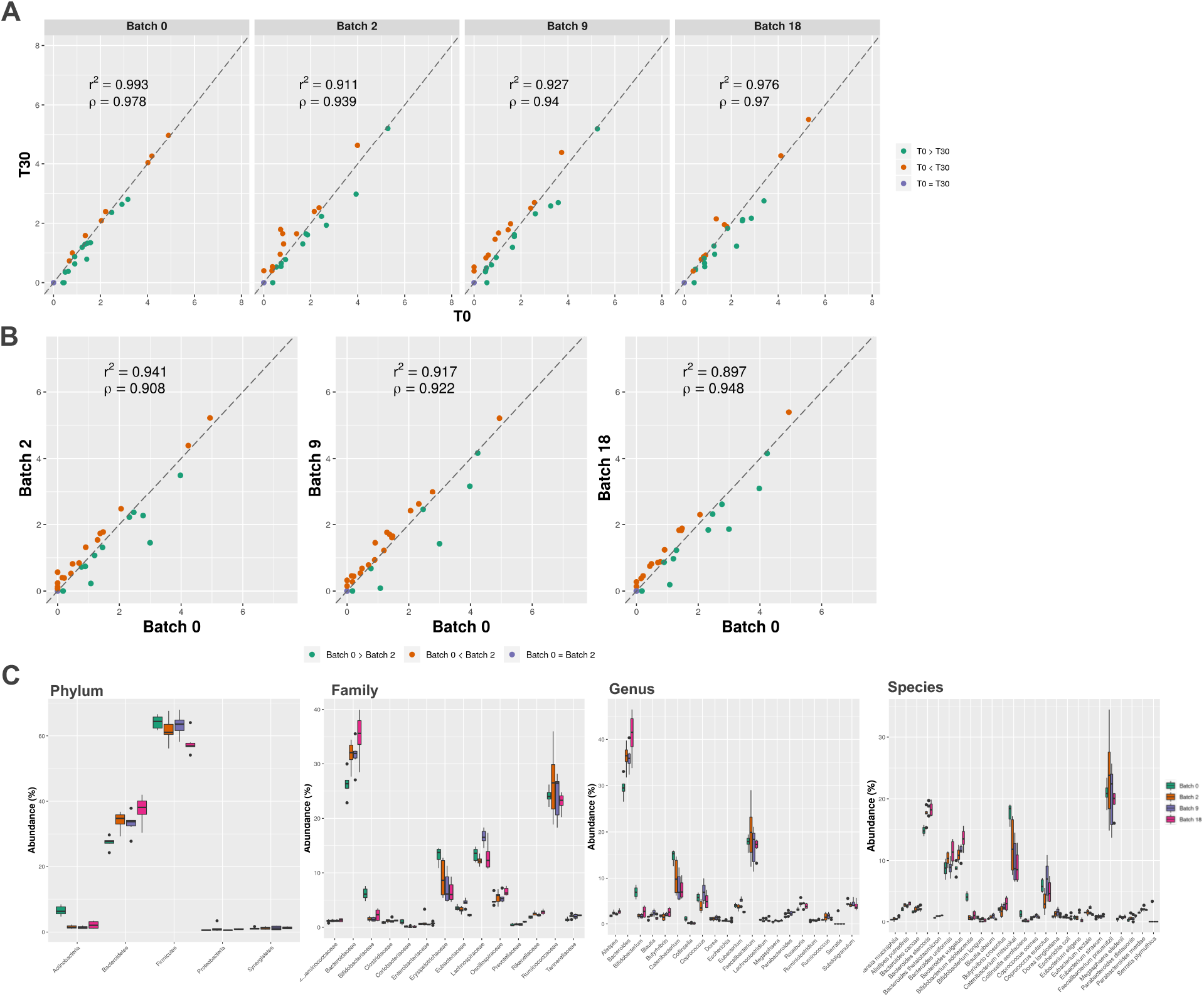
ZSample batch effects in fecal sample storage. Four ZSample lot production (0 - produced in the processing day -, 2, 9 and 18 months of difference from the manufacturing time) were evaluated. **(A)** A correlation analysis showed equivalent results (Pearson - r^2^ and Spearman - ρ coefficients) for the lot solution batches along the time of storage T0-T30, and **(B)** among different lot production batches. **(C)** Bacterial abundances analyzed for phylum, family, genus and species in each production lot. The relative abundance levels are maintained regardless of the solution batch. Lot variations are in the same scale as the intra-replicates variations.

In addition to sample storage, DNA extraction from fecal samples in ZSample was evaluated for four different methods (Supplementary Figure 3). We observed higher correlations and similar diversities for the bacterial profiles obtained with DNeasy PowerSoil, DNeasy PowerSoil Pro modified and QIAamp PowerFecal DNA kit. The recently launched DNeasy PowerSoil Pro kit recovers a higher amount of DNA on average, showing an increased abundance of Firmicutes with reduced Bacteroidetes, Proteobacteria and Verrucomicrobia (Supplementary figure 3). Moreover, no differences related to ZSample solution in the different methods of DNA extraction were observed.

### High throughput amplicon sequencing robustness and analysis using EncodeTools Metabarcode pipeline

Even with the possible variations intrinsic of the method and process, the 16S rRNA amplicon approach must be highly reproducible. Based on this, we performed repeatability and reproducibility (robustness) tests to evaluate our method bias and variations in amplicon library preparation along with the bioinformatics analysis. For the intra-run technical reproducibility test (Supplementary Figures 4A-C) compared to the inter-run reproducibility tests (Supplementary Figures 4D-F) high correlations were obtained, with lower variations in samples alpha and beta-diversities. The overall within-sample correlations for the results obtained with the three different operators can be seen in Figure 3A. Considering all the library and sequencing process variation, different operators, reagents’ lots, plastics and laboratory equipment (*e.g.* thermocyclers and pipettes) the Pearson and Spearman correlation indices showed considerably high values, mainly above 0.9 for all samples. Alpha diversity for the three independent batches of amplicon library preparations and sequencing, performed by three different operators, showed equivalent indexes (Figure 3B). Beta diversity analysis also demonstrated sample-related grouping patterns, which indicates within-sample distances were consistently smaller than between-sample distances (Figure 3C). Negative controls showed a small number of sequenced reads (from 10 to 45), with different and random profiles, while the samples themselves presented from 1,882 to 47,528 reads with consistent bacterial pattern among the replicates.

**Figure 3.**
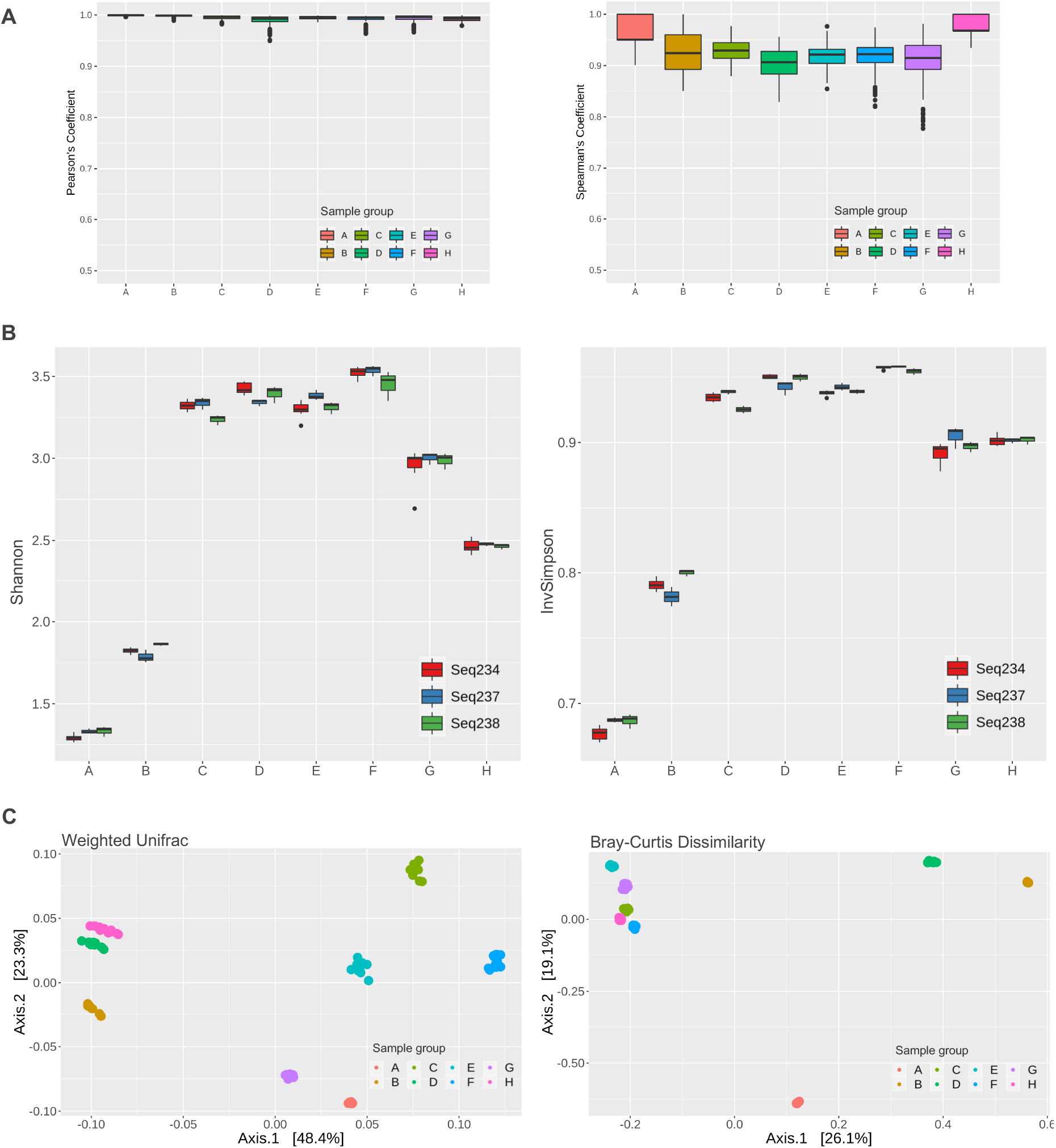
Method reproducibility for DNA library preparation, sequencing and analysis. Eight different DNA sample libraries (A-H) processed in replicates by three different operators and sequenced in three different sequencing runs (Seq234, Seq237, Seq238) all of which analyzed with the EncodeTool metabarcode pipeline. (A) Intra-replicates variations were assessed through correlation analysis demonstrating satisfactory results, with Pearson >0.9 and Spearman >0.8. (B) Alpha diversity indexes, Shannon and InvSimpson, obtained for replicates in each sample set were compared in parallel, showing very small differences throughout the results. (C) Beta-diversity analysis using weighted UniFrac and Bray-curtis dissimilarity, showed that each sample bacterial profile remains clustered together, confirming that variations observed in replicates are less relevant than the original bacterial composition from the different samples.

All the analyses presented in this paper were performed using the EncodeTools Metabarcode pipeline, as described in methods. Besides providing more reliable taxonomic classification due to the LCA feature, this pipeline allows us to access the oligotypes present in a given sample that corresponds to the real amplicon sequence variants (ASVs), and are independent of taxonomic assignment. Oligotype information provides a higher-resolution view of the sample diversity and its DNA sequence composition, so we used that approach to evaluate both our pipeline (EncodeTools Metabarcode) and the robustness of our amplicon library preparation method. As observed in Figure 3 and Supplementary Figure 4, satisfactory correlations and small within-sample variability were observed for the conjunction of experimental methods and the bioinformatics pipeline. To further characterize the latter in terms of sensitivity and specificity, we also extended the analysis to a bacterial mock as described below.

Specificity and sensibility of the EncodeTools Metabarcode pipeline were measured using the bacterial mock results (sample - H) along with the robustness assays. 88.2 ± 2.7% sensitivity and 100% specificity for species level was achieved, given the possible resolution of taxonomical assignment for some 16S rRNA sequences (Supplementary Figure 5). Meanwhile, 99.3 ± 2.7% sensibility and 100% specificity was achieved for the genus level. At family level, the sensibility and specificity reached 100%.

The EncodeTools Metabarcode pipeline generates as output an out_metabarcode (Supplementary Table 2). In this table, we can verify all the oligotypes identified in the analysis, the total number of reads for each oligotype, and the taxonomic assignment given to each oligotype - along with their assigned taxonomic lineage (kingdom, phylum, class, order, family, genus and species). This lineage path stops at the last level in which the oligotype could be classified. For example, several Enterobacteria can only be classified at the family level due to the high similarity among their 16S rRNA gene sequences. When the EncodeTools pipeline matches an oligotype with two or more identical reference sequences, belonging to different species, genus or other higher taxonomy level, the oligotype taxonomic assignment is set for the last common level (ancestor) in the taxonomic path. For instance, if an oligotype could not be resolved at the species level, giving its sequence similarity with two or more species, it probably will be classified at the genus or family level. The out_metabarcode table shows us what are these taxonomies, their identities, and their similarities in the analysis. The read sequence for each oligotype can also be visualized in this table, along with a list of samples in which that given exact sequence was found.

### Brazilian bacteriome profile

The experimental procedures and analyses evaluated in this paper were applied to a subset of over 200 random fecal samples from the Brazilian population. A total of 8,654,114 reads were obtained with an average of 42,010 reads by sample and 2,080 unique oligotypes ranging from 10 to 451,065 reads in the global result. The number of bacterial oligotypes for each sample varied mostly between 30 to 90 (Figure 4A) well approximating a Gaussian distribution (Shapiro-Wilk, P = 0.596) in the populational subset evaluated. On average, taxonomic assignment through the EncodeTools Metabarcode pipeline could be obtained for 98.93% of the reads at the bacterial kingdom level, 97.25% at phylum, 91.82% at family, 81.85% at genus, and 59.35% at the species level (Figure 4B). In this sample subset, phylum, family and genus distributions did not present a Gaussian pattern (Shapiro-Wilk, P <0.01) (Figures 4C, 4D, 4E and 4F) while species are more normally distributed (Shapiro-Wilk, P= 0.145).

**Figure 4.**
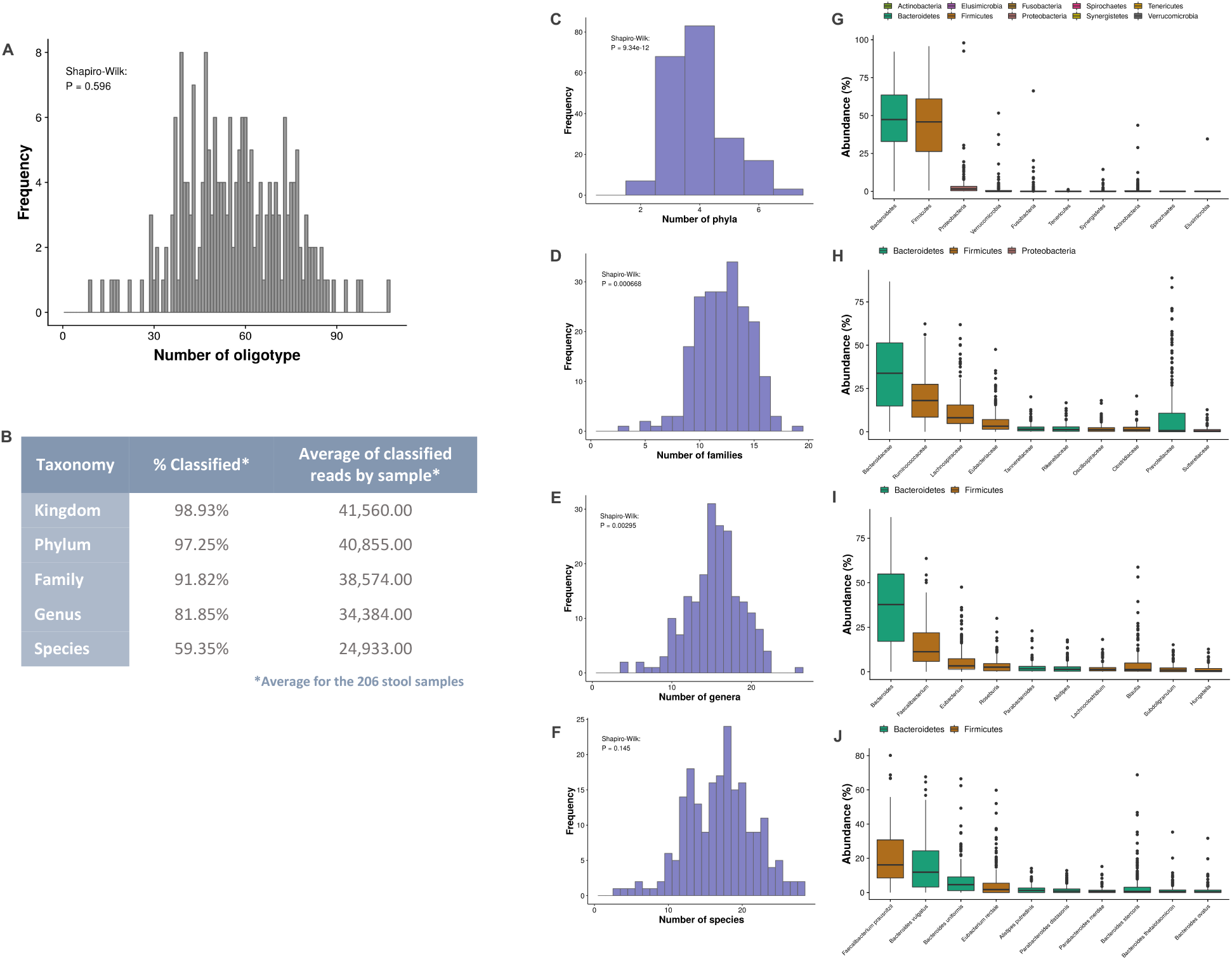
Bacterial profile for the Brazilian fecal microbiome. Over 200 fecal samples from a mischaracterized population were evaluated. **(A)** EncodeTools Metabarcode pipeline showed an oligotype frequency distribution along the samples with more frequent values among 30-90 oligotypes by subject. **(B)** Each sequenced sample yielded approximately 41,500 reads and the EncodeTools Metabarcode Taxonomy Assignment algorithm could attribute phylum taxonomy for an average of 98.93% sample reads. 81.85% of the sequenced reads could be identified at genus level and 59.35% at species level, representing an average of 24,933 reads classified. **(C-F)** Populational distribution of taxonomic assignments in phylum, family, genus and species. Most frequently a subject has between 3 and 4 bacterial phyla, 10 to 15 bacterial families, 10 to 20 bacterial genera, and 12 to 22 bacterial species. Most abundant bacteria for each taxonomy level **(G)** phylum, **(H)**, family, **(I)** genus and **(J)** species are shown accordingly with their populational median distribution.

Regarding the taxonomic assignment, Bacteroidetes and Firmicutes are the most abundant phyla detected in the Brazilian samples with a median abundance values near to 50% (Figure 4G), followed by phyla Proteobacteria, Verrucomicrobia or Actinobacteria, being the lasts, detected in much lower abundances. In consequence, the most abundant families, genera and species are dominated by taxonomies from Bacteroidetes and Firmicutes phyla.

Families Bacteroidaceae, Ruminococcaceae, Lachnospiraceae and Eubacteriaceae are the most abundant families (Figure 4H). Prevotellaceae is also abundant, though its distribution showed relatively lower median and strong positive-skewness, *i.e*., many high-abundance outliers. *Bacteroides* was the most abundant genera detected, followed by *Faecalibacterium*, *Eubacteria* and *Roseburia* (Figure 4I). At the species level, considering the taxonomies that could be reliably resolved by the EncodeTools pipeline (reflecting ~59.35% of the sequenced reads), *Faecalibacterium prausnitzii* is the most abundant species detected in this sample subset (Figure 4J), followed by *Bacteroides vulgatus*, *Bacteroides uniformis*, *Eubacterium rectale* and *Allistipes putrenidis*. Large amounts of *Bacteroides* could not be classified at the species level.

Diversity analysis were performed to visualize how these bacterial profiles are distributed in the populational subset evaluated. Alpha diversity indexes (Chao1, Shannon, Simpson and InvSimpson) were calculated for the samples oligotypes (Figure 5A). Chao1 was the only index with a normal distribution (Shapiro-Wilk, P= 0.596). Other indexes did not show a Gaussian distribution; however, they are skewed for some common ranges. The same alpha diversity analysis was performed for phylum, family, genus and species (Supplementary Figures 6A-D). However, all of them presented lower diversity indexes, as expected due to oligotype clustering in higher taxonomic ranks, reducing the number of taxonomies to account for the analysis. None of these presented Gaussian distribution, except for Chao1 at species level (Supplementary Figure 6D).

**Figure 5.**
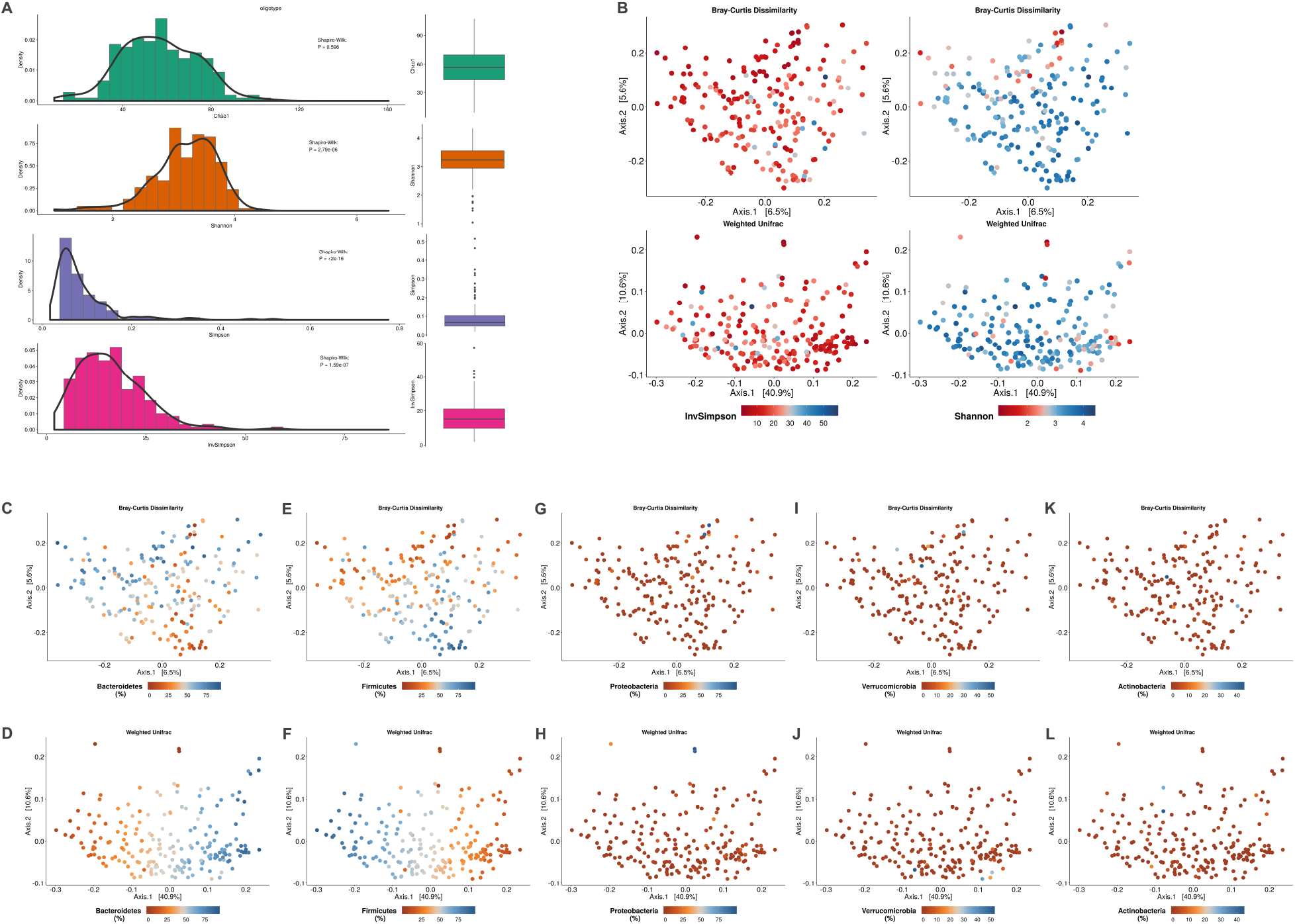
Brazilian bacteriome diversity analysis with oligotype sequences. **(A)** Alpha-diversity Chao1, Shannon, Simpson and InvSimpson indexes distribution for the data were analyzed. Only Chao1 index approximates to a Gaussian distribution (Shapiro-Wilk 0.59), however other alpha diversity indexes showed pretty narrow skewed data distribution. **(B)** PCoA plots for Bray-Curtis and weighted UniFrac showed a lack of correlation regarding alpha-diversity Shannon and InvSimpson distributions. **(C-L)** Bray-Curtis dissimilarity and weighted UniFrac PCoA plots colored by phylum abundance distribution. Most abundant phyla Bacteroidetes **(C-D)** and Firmicutes **(E-F)** have marked distributions among all the samples analyzed. Three major groups could be seen, samples higher in Firmicutes (that are lower in Bacteroidetes), samples lower in Bacteroidetes (higher in Firmicutes) and samples with equivalent amounts of both phyla. Other less abundant phyla **(G-H)** Proteobacteria, **(I-J)** Verrucomicrobia and **(K-L)** Actinobacteria doesn’t seem to contribute to the populational distribution observed in the beta-diversity analysis.

The beta diversity analyses, using both Bray-Curtis dissimilarity and phylogenetic similarity Weighted UniFrac, were performed for the samples’ oligotypes. PCoA plots showed that samples were widely dispersed, without specific sample subgroups. In Figure 5B, alpha diversity indexes (Shannon and InvSimpson) didn’t seems to explain any significant pattern of sample distribution among the population subset. However, the PCoA arrangement, considering the first two Principal Coordinates (PC’s) for both methods, seems to be guided by the two most abundant phyla in the samples: Bacteroidetes and Firmicutes. Samples with higher abundance of Bacteroidetes have smaller amounts of Firmicutes and samples with less Bacteroidetes have more abundant oligotypes attributed to Firmicutes (Figures 5C-F). Less abundant phyla have more homogeneous low abundance distribution among samples (Figures 5G-L). Lower taxonomic levels (family, genus and species) seem to be less correlated with the overall beta-diversity arrangements, at least when considering the first two PCs. Still, some small groupings can be observed for samples with higher amounts of Bacteroidaceae, Ruminococcaceae, Prevotellaceae, *Bacteroides*, *Faecalibacterium*, *Faecalibacterium prausnitzii* and *Bacteroides vulgatus* (Supplementary Figure 6). Other abundance-driven grouping tendencies can be observed in the PCoAs shown in Supplementary Figure 6. However, further analyses are required to establish specific correlations between certain taxonomies and possible enterotypes/bacteriome profiles.

Finally, 30 negative controls of DNA extraction (CNE) were analyzed along with 44 negative PCR reaction controls (CNR). This control analysis, performed at each sequencing batch, allows us to detect deviations in the process that could invalidate the sample results. Here, the oligotype numbers as well as the total reads per library obtained for control samples were notably low, which yielded completely different alpha and beta diversity profiles (Supplementary Figure 7). Thus, no significant batch contamination from reagents was detected and the process is capable of reliably representing the original bacteriome samples’ compositions.

## Discussion

In this paper, we present an end-to-end assessment of the methodologies that we developed to analyze the bacterial composition of the intestinal microbiome. First, we created a sample collection kit, Probiome, that people can easily take home and use to collect a small amount of fecal sample with a sterile swab and store it at room temperature using a tube containing a stabilizing solution to deliver it to the laboratory within 30 days after sample collection. Then the laboratory performed the following procedures: DNA sample extraction and 16S rRNA amplicon sequencing to access the sample bacterial composition through a bioinformatics - EncodeTools Metabarcode pipeline. All these processes were evaluated to account for processing variabilities and reproducibility of the obtained results.

A very high load of microorganisms populates our gut. From the moment of sample collection to the DNA extraction, the bacterial profile can suffer dramatic changes caused by sample degradation or even microorganisms overgrowth. It may favor the detection of some microorganisms over another (*e.g.*, aerobes x anaerobes). Thus, adequate sample storage is necessary until proceeding to the DNA extraction to preserve the real bacterial profile in the samples [14,15,36]. Although immediately freezing seems to be the best choice [36] it is not feasible in large-scale populational studies. Some storage solutions have already been evaluated like RNAlater, OMINIgene-gut, Norgen, Shield, Tris-EDTA, ethanol 70%, 90% or 95% and FTA cards [14,15,36–38]. Generally, these studies evaluated the sample preservation at the short term, from two to seven days, and reported that OMNI, ethanol 95%, Norgen and FTA cards were the best preservation alternatives. However, only OMNI and Norgen were shown to impairs bacterial growth in the sample, while RNAlater should be avoided given its poor DNA recovery and alterations in bacterial taxa recovered [14,36,38–40]. Only one of these studies performed a long-term survey of sample preservation at room temperature for eight weeks, showing that OMNIgene-gut, FTA cards and ethanol 95% were the best preservatives with very minimal variations, comparable to technical replicates variations [14]. Another long-term study evaluated 5-year samples stored in RNAlater and frozen at −80 °C. However, these samples remained 6-17 days at room temperature before freezing [41] so this study did not account for the alterations in the microbial profile caused by the room temperature storage during a considerably long period - which is critical given the previous research warnings to avoid RNAlater. Based on all this knowledge, together with the high costs of solutions like OMNI or Norgen and the need for an accessible fecal collection kit in Brazil, we developed our storage solution, ZSample. It was tested regarding bacterial inactivation and profile maintenance for 30 days at room temperature. Variations in the bacterial profile related to different lot productions of the solution were not detected either.

After sample collection, the storage lasts until the DNA extraction process, which obtain the microbial genetic information of the sample. This is also an intensive subject of investigation, since different DNA extraction methods can lead to different microbial profiles. Even though subject’s differences are known to be one of the greatest sources of variability for human microbiome data, some DNA extraction methods yield more variations than others [16,42–45]. We detected significant variations in the bacterial profile recovered by DNeasy PowerSoil Pro kit. These variations may be attributed to the bead-beating with the zirconium beads during the lysis process. To the best of our knowledge, DNeasy PowerSoil and QIAamp PowerFecal DNA represent the most used kits in microbiome research studies. Hence, aiming to keep consistency with the microbiome profiles reported in the literature, we continued the use of PowerSoil Kit. Nonetheless, it remains to be confirmed which DNA extraction kit yields the most reliable results, *i.e.,* the one which most closely resembles the original samples’ bacterial composition.

Besides sample storage and DNA extraction, the DNA library preparation for high-throughput sequencing could also have a greater impact on the assessment of the results. In general, there are two main approaches used to assess the gut microbial diversity: a metabarcode analysis, such as 16S rRNA gene for bacterial identification and compositional analysis, and metagenomics approaches which, in addition to bacterial identification, can reveal other microorganisms such as fungi, viruses or eukaryotes, as well as their interaction networks through genes and metabolism inferences. Both methodologies are valid and should be applied in accordance with the expected results. To perform a high-level community profiling, 16S rRNA marker gene is most indicated, whereas to perform functional profiling, metagenomics must be used [13]. Additionally, previous research as the MetaHIT project demonstrated that human intestinal microbiome is composed mainly of bacteria, more than 90% of the intestinal DNA recovered was bacterial-related [2]. Also, it was shown that 16S rRNA amplicon sequencing recovers more bacterial diversity than shotgun based metagenomics [46]. Thus, 16S rRNA marker gene amplicons are best suited for our analysis and expected results, being the method of choice for this study.

We evaluated the reproducibility of our amplicon library preparation performing several replicates and including variables such as different operators, equipment, reagents and dates of processing. These assays were also used to test our pipeline of analysis (EncodeTools Metabarcode) justifying the higher number of replicates performed and a bacterial mock sample with known composition. The EncodeTools Metabarcode pipeline was developed to provide more reliable results, assessing single variations from the sequences with greater confidence and improved taxonomic assignment.

The analysis of amplicon sequencing variants (ASVs), grouped into oligotypes composed by sequences with 100% similarity, is the main feature that improves the 16S rRNA gene bacterial profiling [13,17]. We already use this approach of reads clustering since 2014, for hospital microbiome surveillances using 16S rRNA gene high-throughput sequencing [26]. Currently, new bioinformatics tools are available to assist in the accuracy of obtained sequence reads, as the denoising procedures based on software packages like Deblur and DADA2 [18,22,47]. These pipelines help in the detection of sequencing artifacts and erroneous reads, giving a more reliable result regarding oligotypes that may vary by only one nucleotide, as well as being more useful in the detection of real variations among samples [22].

In the EncodeTools Metabarcode pipeline, we implemented a *de novo* taxonomic assignment, based on similarity [25], which can classify most of oligotypes at least to the phylum level. Thus, associating our EncodeTools metabarcode pipeline with the 283bp - 16S rRNA V3/V4 oligotypes and a *de novo* taxonomic classification, we can obtain high-quality, highly-reproducible results. Regarding taxonomic assignment, using a read length of 283bp provides a great improvement for taxonomy resolution at several ranks, including at species level. This approach seems to perform even better than some metagenomics approaches in which only 52.8% of the fragments could be assigned to genus and 80% to phylum - while still reporting bacterial dominance within intestinal microbiome [2].

The inclusion of negative controls along the process is also important to assess possible contaminations that may occur in the DNA extraction, PCR amplification, sequencing or even in the bioinformatics pipeline, as previously reported [16,48,49]. Contaminations with some bacterial DNA is ubiquitous among DNA extraction kits and laboratory reagents [48], being more relevant for microbial detection in low-biomass samples [50]. In general, we detected very low number of reads in negative controls, with an average of only four oligotypes and highly random bacterial profiles. In each experimental batch, these contaminations must be evaluated to understand the magnitude of their impact on the results, whether they can be filtered from some samples or even if they invalidate the entire result. EncodeTools Metabarcode pipeline has this filtering options embedded in its code to evaluate negative controls from each experimental batch.

The procedures described here were applied to a batch of more than 200 fecal samples collected from the Brazilian population. So far, there were no reports for the microbial diversity of the Brazilian gut microbiome, thus we presented a first general overview of this profile for bacterial abundance and distribution. The Brazilian fecal microbiome samples have a consistent distribution of oligotypes, phylum, family, genus and species along the population analyzed, often approximating Gaussian distributions. Alpha and beta diversities have similar distributions to those reported by other studies [51], in which the main populational dissimilarities are guided by the most abundant phyla. Generally, most of the studies published so far showed that the human intestinal microbiome is mainly composed by Bacteroidetes and Firmicutes phyla [2,3,5–7,9,52], as we observed here. The Brazilian microbiome profile shown here should be further investigated with stratified metadata to better understand patterns and microbial diversities related to populational geography, diet, age, sex, and several other possibly associated/confounding factors. Brazilians compose a very diverse and geographically distributed population. Deep characterization of their microbiome profiles is necessary if we want to better comprehend the applicability of the information derived from other populational studies [8,12,53].

In conclusion, we provided an end-to-end assessment of microbiome sample processing and analysis, as well as its applications to the study of the Brazilian fecal bacteriome. Using an effective sample collection method, with a standardized sample processing, DNA sequencing and bioinformatics analysis, we achieved highly reliable results. One of the major gains of the methodology herein presented is the bioinformatics pipeline in which oligotypes represent the pure sample diversity, free of biased taxonomic assignment for generalist groupings as in OTU picking [13,22,54,55]. OTUs are known to underestimate sample diversity. However, sOTUs, ASVs or oligotypes approaches overcome this issue, and even empower 16S rRNA studies to reveal more bacterial diversity than shotgun metagenomics [46,56]. In addition to the oligotype approach, we also gain phylogenetic resolution by sequencing a larger fragment that most studies do, which improves taxonomical assignment in fecal sample characterizations. Using these methodologies, larger sample cohorts should be analyzed for Brazilian population and more detailed comparative studies and meta-analyses must increase the knowledge about the intestinal microbiome of such a diverse population.

## Conflict of interest

All authors are or were currently full-time employees of BiomeHub (SC, Brazil), a research and consulting company specialized in microbiome technologies.

## Funding Statement

BiomeHub funded this study.

