## Supplementary figures for "End-to-end assessment of fecal bacteriome analysis: from sample processing to DNA sequencing and bioinformatics results"

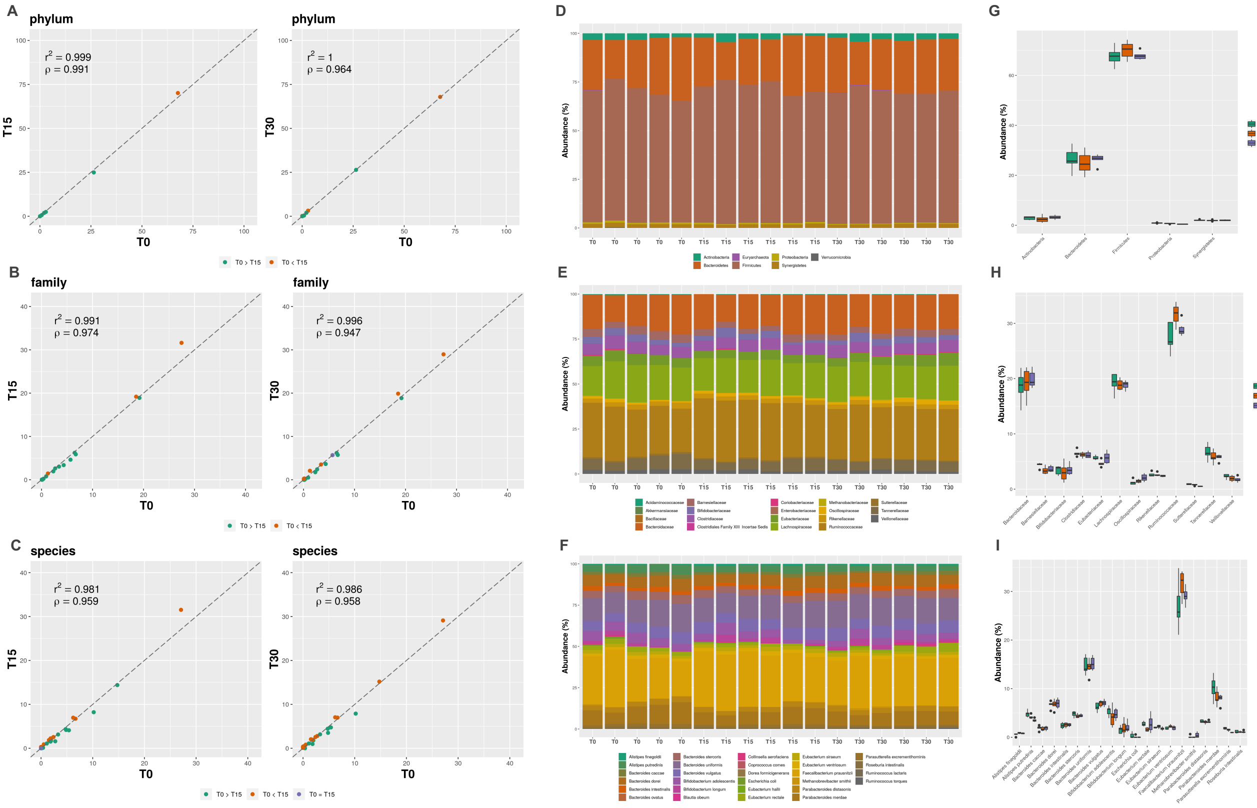


**Supplementary figure 1**. **Fecal sample storage and bacterial profile along 30 days.** Correlation analysis for T0-T15 and T0-T30 obtained results considering data with taxonomical assignment for **(A)** phylum, **(B)** family and **(C)** species. Pearson ($r^2$) and Spearman (ρ) correlations were calculated. Relative abundance bacterial profiles for each sample along with the 30 days storage, including sample replicates for **(D)** phylum, **(E)** family and **(F)** species. Boxplots showing abundances distributions and deviations in each storage time (T0, T15 and T30) for taxonomic levels of **(G)** phylum, **(H)** family and **(I)** species.

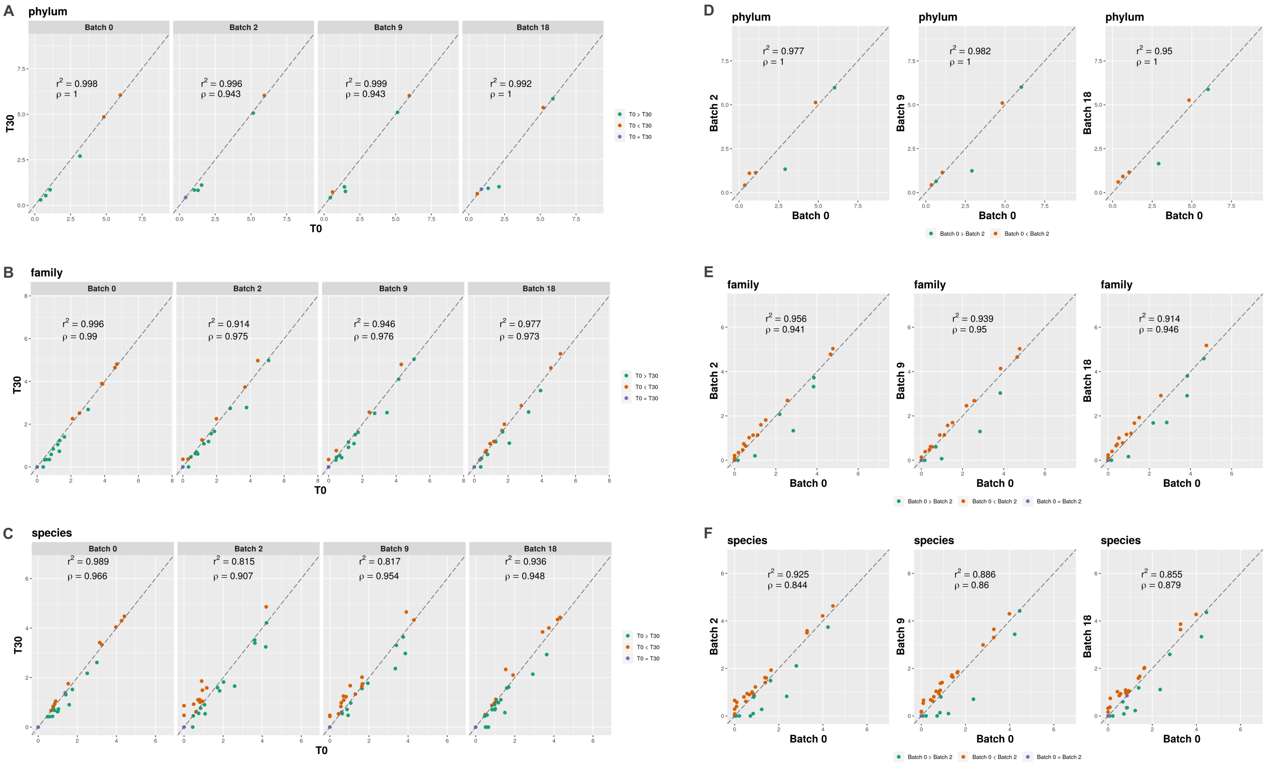


**Supplementary figure 2. ZSample batch effects assessment along 30 days of fecal sample storage.** Correlations (Pearson ($r^2$) and Spearman (ρ) for bacterial profile before and after the 30 days storage in different batches of ZSample lot production. Results are shown for taxonomic assignments of **(A)** phylum, **(B)** family and **(C)** species. Also, correlations among different batches were performed for the same taxonomic levels **(D)** phylum, **(E)** family and **(F)** species. High correlation values $r^2$ and ρ were obtained among the compared results.

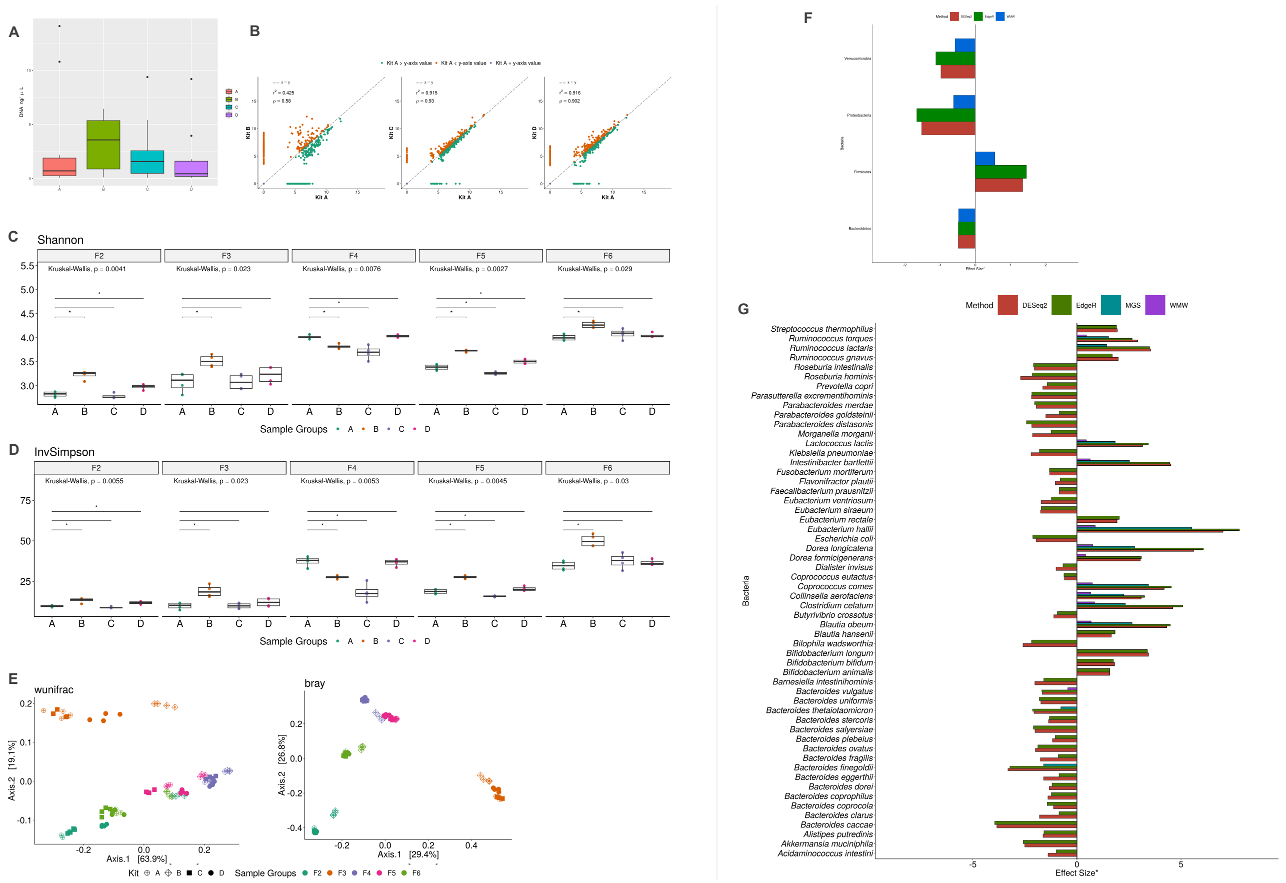


**Supplementary figure 3. Fecal DNA extraction results assessment for different kits.** Four different extraction kits were evaluated A-DNeasy PowerSoil; B- DNeasy PowerSoil PRO; C-DNeasy PowerSoil PRO modified (zirconium beads changed to silica beads) and D-DNeasy Power Fecal. **(A)** Amounts of DNA extracted (ng/ul) were quantified using Picogreen (Invitrogen, USA). Kit B presented the higher overall DNA amounts recovered. **(B)** An analysis correlation performed after bacterial 16S rRNA gene sequencing revealed that kit B presents the most different results regarding the sample bacterial composition compared to kit A ($r^2$=0.4 and ρ=0.58). Kits A, C and D have similar and equivalent results ($r^2$ and $\rho > 0.91$). To better evaluate these differences among kits, Shannon **(C)** and **(D)** InvSimpson alpha diversities analysis were performed for each subject separately. All samples from the B kit presented significant differences (*Kruskall-Wallis, Wilcoxon* p <0.05), generally showing higher alpha-diversity indexes. **(E)** Despite kits variations, beta-diversity analysis (weighted UniFrac and Bray-Curtis) showed that the bacterial profile within an individual is much more consistent than the method of extraction. However, it is clearly visible the deviations resulting from fecal DNA extractions with B kit. Differential abundance analysis with DESeq2, EdgeR, MGS and WMW were performed to identify which are the bacteria phylum **(F)** and species **(G)** deviating between kit A and kit B. It was observed an increase of the phylum Firmicutes and a reduction for Bacteroidetes, Proteobacteria and Verrucomicrobia for kit B. Also, most of the bacteria associated with these phyla were affected and detected at least by two differential abundance methods used. *Effect sizes are fold-changes in log2scale, except for WMW which shows $Z_{score}/\sqrt{N}$.

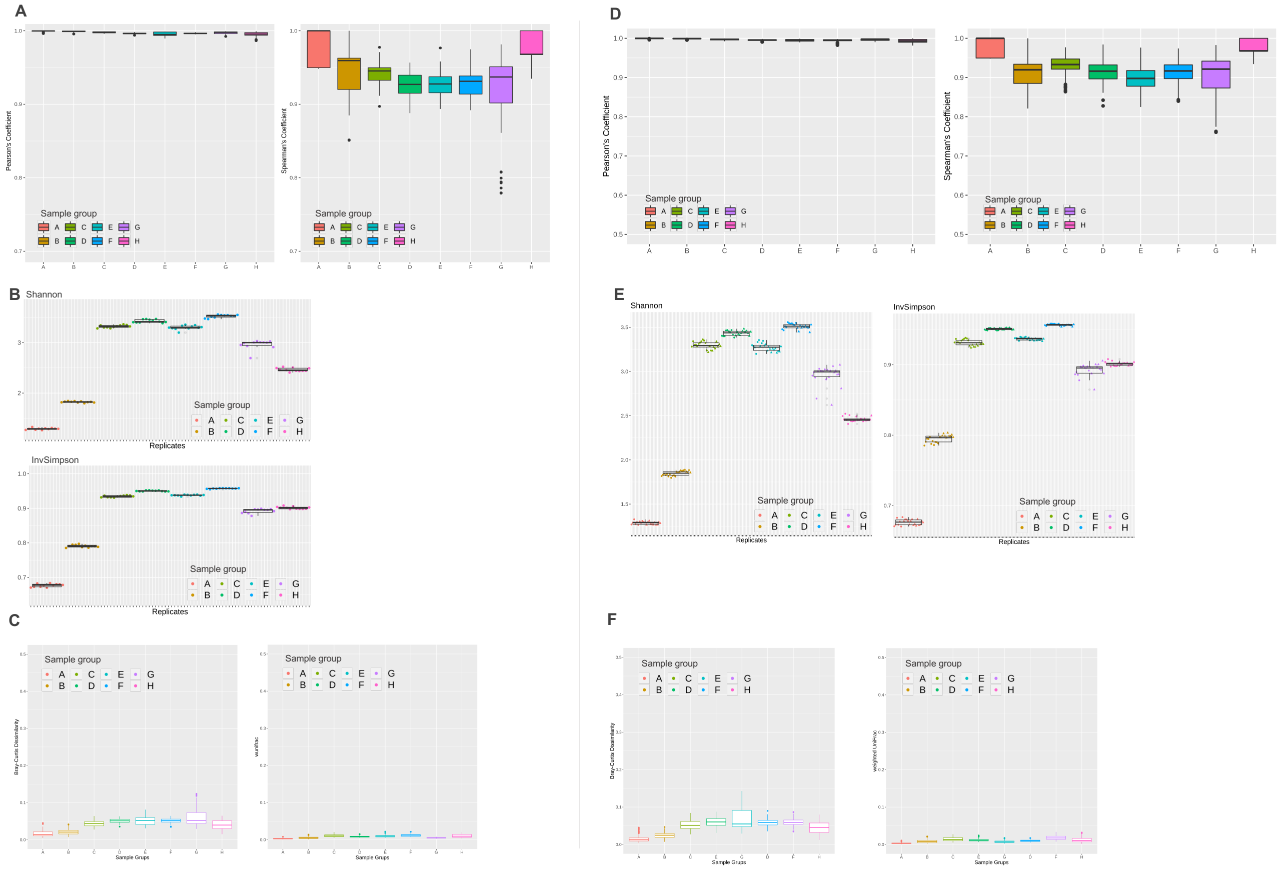


**Supplementary figure 4. Experimental reproducibility for amplicon DNA library preparation, sequencing and EncodeTools Metabarcode analysis.** Different samples subsets for correlation, alpha and beta diversity analysis. **(A-C)** - Subset of 11 replicates performed by only one operator, in a single sequencing run. **(D-F)** - Subset of 11 replicates re-sequenced in a second run and analyzed along with the first sequencing data.

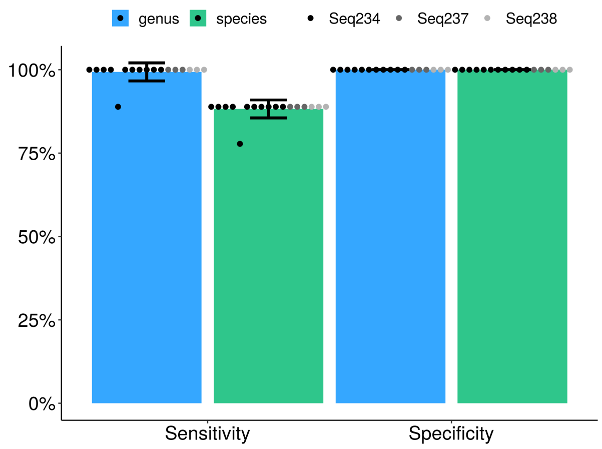


**Supplementary figure 5. Sensitivity and Specificity results achieved for library preparation, DNA sequencing and EncodeTools Metabarcode pipeline.** Sensitivity was evaluated for the ability to recover the expected bacteria and specificity as the confidence level in bacterial detection among a diverse microbial subset. A bacterial mock sample composed of *Acinetobacter baumanii, Bacillus subtilis, Enterococcus faecalis, Escherichia coli, Klebsiella pneumoniae, Listeria monocytogenes, Pseudomonas aeruginosa, Salmonella enterica* and *Staphylococcus aureus* was used. Results obtained for 17 replicates performed by three different operators in three different sequencing runs. Values achieved were 100% specificity at genus and species level, 99.3 ± 2.7% sensibility at genus level and 88.2 ± 2.7% at the species level. At the family level, specificity and sensibility was 100%. These variations occurred mainly because *Listeria monocytogenes* 16S rRNA sequences do not have phylogenetic resolution enough to EncodeTools Taxonomy Assignment algorithm classify them at species. *Listeria* only has resolution to be classified at genus level. Additionally, some sequences from the Enterobacterias do not have resolution to lower classification than family, this occurs mainly for *Salmonella* sequences. Thus, sensibility variations are attributed to the lack of taxonomical resolution in the 16S rRNA sequences evaluated. Also, a deviation observed is due to one replicate with the lowest reads sequencing coverage (2,509 reads).

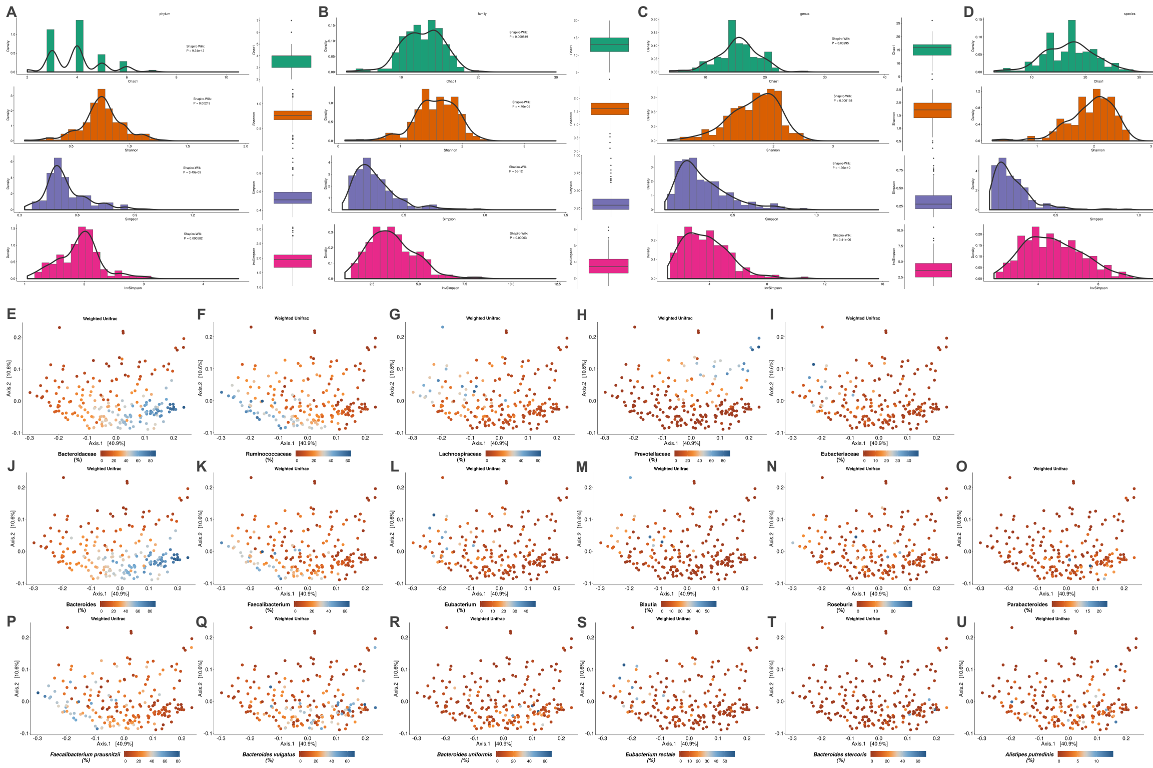

**Supplementary figure 6. Brazilian bacteriome diversities and distributions.** Besides the oligotype alpha-diversity profiles for the 206 Brazilian fecal samples presented in the main text, we also generate Chao1, Shannon, Simpson and InvSimpson indexes based in the EncodeTools taxonomic assignments for **(A)** phylum, **(B)** family, **(C)** genus and **(D)** species. Results distributions were equivalent to oligotypes, however considering these taxonomic ranks the diversity indexes decrease, given the reduced variables after taxonomic assignment. Weighted UniFrac PCoA plots with populational distributions were shown **(E-I)** family, **(J-O)** genus and **(P-U)** species most abundant in the Brazilian dataset evaluated. Bacteroidaceae and Ruminococcaceae families have similar patterns related to their phyla as well as Bacteroides and *Faecalibacterium* genus, also reflecting in *Faecalibacterium prausnitzii, Eubacterium rectale, Bacteroides vulgatus* and *Bacteroides uniformis* species. Prevotellaceae family seems to have a particular grouping for samples with higher abundance of this family that should be further investigated.

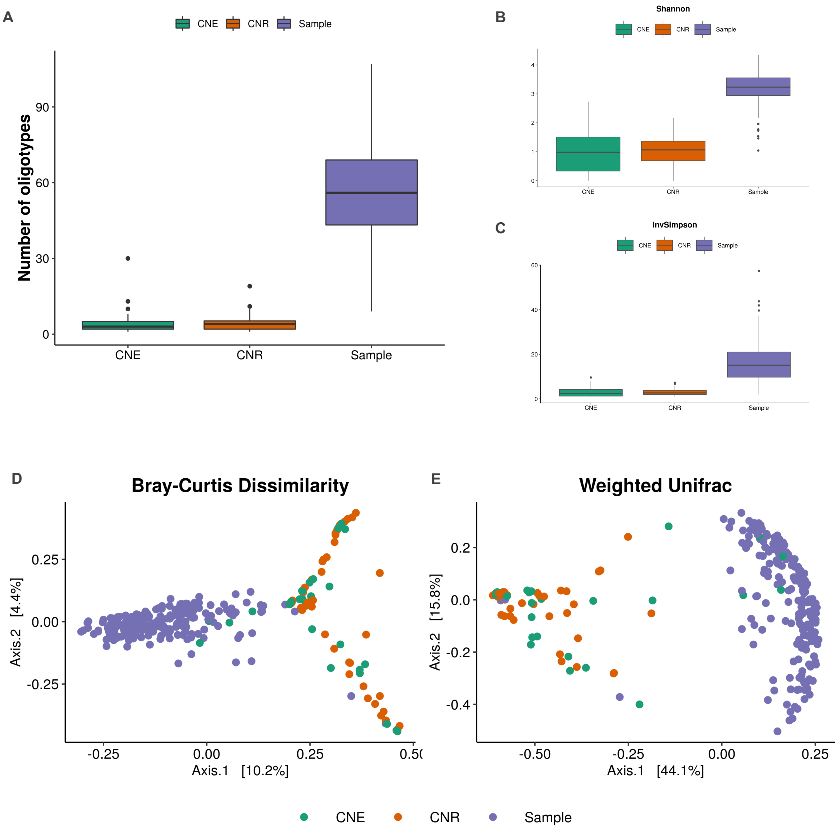


**Supplementary figure 7. Negative controls analysis.** 30 DNA extraction negative controls (CNE) and 44 PCR reaction negative controls (CNR) were analyzed along with the 206 Brazilian fecal samples. **(A)** After EncodeTools Metabarcode analysis it can be observed that negative controls have a much lower amount of oligotypes detected in relation to the samples. **(B-C)** Also, Alpha diversities are much different with lower diversities observed in negative controls, corroborating the lower amount of oligotypes detected. Concerning to the negative controls, the bacterial profile beta diversity analysis like Bray-Curtis dissimilarity and Weighted UniFrac showed highly dissimilar profiles for CNEs and CNRs. Except for four CNEs with similar profile to the samples, all the other ones represent low abundance and random amplification profiles from laboratory reagents contaminants that is widely discussed in high-throughput sequencing bacterial sequencing. These four CNEs with sample-related profile were identified as basal contaminations from the samples in controls and used for an internal investigation, process improvement and validation of the obtained results.
